# Label-Free Optical Biopsy Reveals Biomolecular and Morphological Features of Diabetic Kidney Tissue in 2D and 3D

**DOI:** 10.1101/2024.10.27.620507

**Authors:** Anthony A. Fung, Zhi Li, Craig Boote, Petar Markov, Sanjay Jain, Lingyan Shi

## Abstract

Kidney disease, the ninth leading cause of death in the United States, has one of the poorest diagnostic efficiencies of only 10%^1^. Conventional diagnostic methods often rely on light microscopy analysis of 2D fixed tissue sections with limited molecular insight compared to omics studies. Targeting multiple features in a biopsy using molecular or chemical reagents can enhance molecular phenotyping but are limited by overlap of their spatial and chromatic properties, variations in quality of the products, limited multimodal nature and need additional tissue processing. To overcome these limitations and increase the breadth of molecular information available from tissue without an impact on routine diagnostic workup, we implemented label-free imaging modalities including stimulated Raman scattering (SRS) microscopy, second harmonic generation (SHG), and two photon fluorescence (TPF) into a single microscopy setup. We visualized and identified morphological, structural, lipidomic, and metabolic biomarkers of control and diabetic human kidney biopsy samples in 2D and 3D at a subcellular resolution. The label-free biomarkers, including collagen fiber morphology, mesangial-glomerular fractional volume, lipid saturation, redox status, and relative lipid and protein concentrations in the form of Stimulated Raman Histology (SRH), illustrate distinct features in kidney disease tissues not previously appreciated. The same tissue section can be used for routine diagnostic work up thus enhancing the power of cliniopathological insights obtainable without compromising already limited tissue. The additional multimodal biomarkers and metrics are broadly applicable and deepen our understanding of the progression of kidney diseases by integrating lipidomic, fibrotic, and metabolic data.

Graphical Abstract
Label-free indicators of diabetic nephropathies.

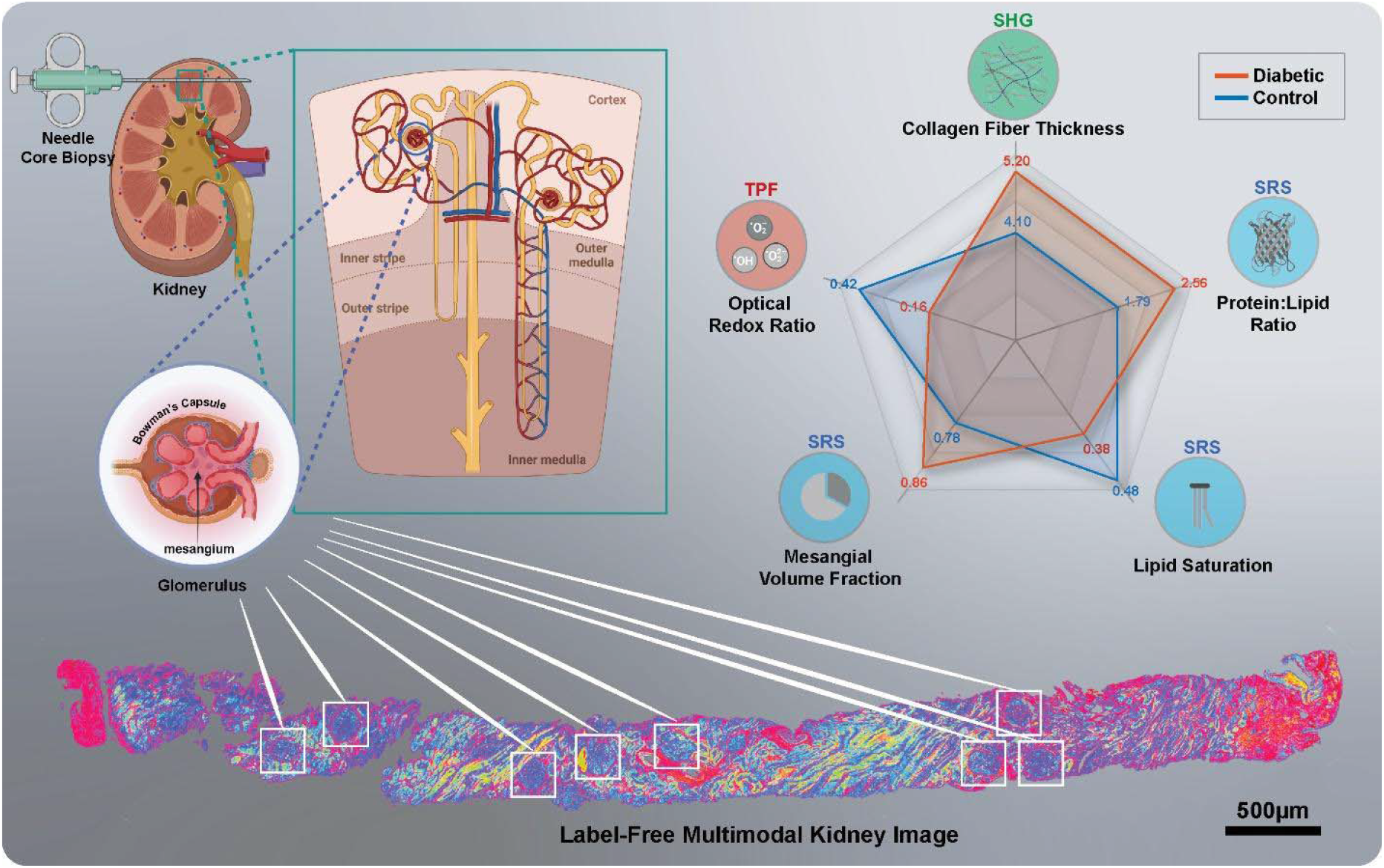

## Introduction

With over 40% of diabetic patients developing diabetic kidney disease (DKD), the leading cause of chronic kidney disease (CKD)^2^, diabetes is one of the most significant risk factors in kidney-related mortality. It can affect renal tissue both morphologically and molecularly, altering redox balance, lipid metabolism, and collagen homeostasis. To afford the highest diagnostic power, multiplexing capability is often a prerequisite when considering any tool or pipeline. Conventional staining methods used for kidney biopsy include Hematoxylin and Eosin (H&E), methenamine silver, Masson’s trichrome staining, and immunofluorescence^3^. In practice, however, multiplexing capabilities are limited by the chroma of the stain or absorption/emission wavelengths of the fluorescent probe. Several washes in acids or alcohols may be required to re-color the sample with a different stain^4,5^, but this process can alter the native lipid distribution^6^ and affect future fluorescent staining. MALDI/MS based methods provide impressive label-free multiplexing capabilities but may entail professional matrix embedding, have limited subcellular spatial resolution, affect tissue integrity and cannot capture true 3D images. To circumvent these issues, adjacent sections of the tissue may be used, but may exhibit different morphologies, suffer from physical slicing deformations, and require larger biopsies from the patient putting constraints on already limited tissue. Studies have reported distinct glomerular^7^, tubulointerstitial^2,8^, and medullary^9^ changes in DKD. Therefore, there is a demand for whole slide imaging (WSI) to maintain the spatial context of any biomarkers.

Morphological changes in DKD include an inflamed mesangial fractional volume [Vv(Mes/Glom)], defined herein as the fraction of glomerular volume occupied by tuft (including mesangial cells, capillary cells, and podocytes), and thicker collagen fibers in the glomerular basement membrane, interstitial matrix, and accumulation within the mesangium^10–16^. Studies also show that CKD tissues exhibit a larger Vv(Mes/Glom) due to mesangial expansion^10–13,17^. This phenomenon has also been linked to high density cholesterol dyslipidemia^18–20^. However, this volume ratio is often measured from cross sections of tissue using pixel density, point sample intercept (PSI), or lineal analysis^13,21–23^. In a biopsy, glomerular units are often found in different focal planes, which may complicate diagnoses by reducing the statistical power of the analysis. With sufficient sampling, variations within a patient sample become negligible^24^. Recently, a thorough evaluation of various glomerular estimation methods confirmed that different methods may misrepresent the true glomerular volume^25^, but presently there is no study that directly compares these with 3D imaging results^26–30^. Interestingly, mesangial expansion is relatively slow to develop and is reminiscent of advanced CKD, rather than only for DKD^7,31^. Could this be due to the large variance in Vv(Mes/Glom) measurements? A stimulated Raman scattering (SRS) microscopy image of the CH_3_ stretching region (2940 cm^−1^) shows a human kidney biopsy with glomeruli in various focal planes (**Fig. S1a**). This exemplary image illustrates how polar slices of glomeruli can have starkly contrasted mesangium-to-glomerulus area ratios. Serial slices just 10 µm apart can cause significant differences in this fractional volume (**Fig. S1b**). Therefore, a focus on a more inclusive field of view is critical to fully utilize precious biopsy samples.

Biomolecular alterations in DKD include lipid dysregulation, proteinuria, or altered NAD(P)H and flavin levels. These features are not entirely disjointed from the morphological differences. Fibrogenesis is heavily influenced by endothelial cell function, particularly in peritubular microvessels^32^. In mice, the mitochondrial deacetylase, Sirtuin 3 (SIRT3), is not only a tight regulator of glucose and lipid metabolism, but a key metabolic programmer of renal fibrosis as well^33^. Kidney tissue has a relatively low level of fatty acid synthase, so much of the lipid metabolism is undertaken by the incorporation of exogenous lipids and the mitochondrial-mediated breakdown of fatty acids. Dyslipidemia is frequently observed in nephrotic syndromes and kidney diseases and is often presented as elevated levels of low-density lipoprotein (LDL) cholesterol and decreased levels of high-density lipoprotein (HDL) cholesterol in sera^34^. Oxidative stress has been associated with both fibrosis and lipid peroxidation. As in previous study we used the normalized optical redox ratio (ORR) as an indicator of oxidative stress, which can be measured using the label-free two-photon autofluorescence imaging of NAD(P)H and flavins acquired at 460nm and 525nm, respectively^35^.

In the following work, we leveraged second harmonic generation (SHG), stimulated Raman scattering (SRS), and two photon fluorescence (TPF) in a single microscopy platform (**Fig. 1a-b**) to generate a label free molecular indicators of diabetic kidney tissue pathology without additional usage of precious kidney tissue. Major macromolecular classes such as saturated and unsaturated fatty acids, proteins, collagen, flavins, and NAD(P)H are interrogated with high spatial resolution, and result in 5 major label-free biomarkers spanning the morphological and molecular space; lipid saturation, oxidative stress, protein/lipid ratio, interstitial collagen, and fractional glomerular volume. This platform is capable of multimodal whole sample (**Fig. 1c-f**) or region-specific (**Fig. 1g-h**) imaging and analysis. A representative multimodal image of a control sample can be found in **supplementary figure 2**. Additionally, this platform can be executed in 3D up to a depth of 200 µm non-destructively, maintaining the flexibility to carry out other analyses downstream. In the following results section, we apply the multimodal imaging platform to identify morphological (including mesangial proliferation, collagen fiber anisotropy and thickness) and biochemical (including lipid accumulation, lipid saturation, and optical redox ratio) biomarkers in diabetic tissues, as well as discuss their impacts and significance in the current scientific landscape.

**Figure 1.**
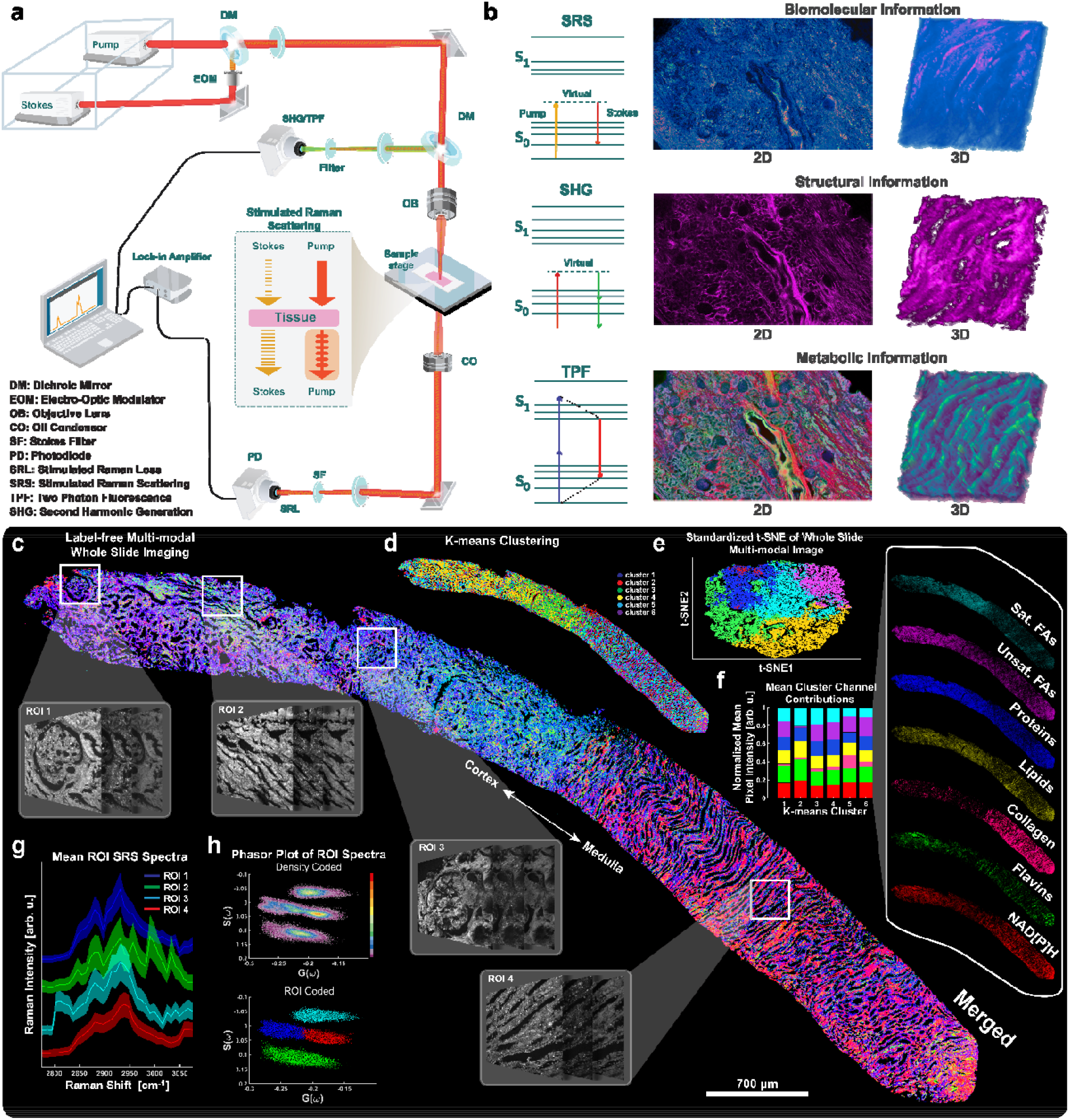
Design and capabilities of the label-free optical platform span targeted regions of interest to whole slide imaging. **a**, Diagram of the label-free microscopy system showing the transmission detection of stimulated Raman loss and epi detection of fluorescence emission and second harmonic generation. **b**, Example images corresponding to each modality in 2D and 3D, along with their respective Jablonski diagram. **c**, A whole slide image of a renal core biopsy from a patient diagnosed with Diabetic Nephropathy (DN) imaged using the platform described in **a**. The image is the result of overlaying seven discrete channels ascribed to saturated fatty acids (SRS at 2880cm^−1^), unsaturated fatty acids (SRS 3011cm^−1^), proteins (SRS 2940cm^−1^), lipids (SRS 2850cm^−1^), collagen (SHG at 1031nm/515nm), flavins (TPF 860nm/515nm), and NAD(P)H (TPF 780nm/460nm). **d**, An unsupervised k-means clustering of the multimodal WSI. **e**, A PCA-initialized, standardized t-SNE plot of the pixels in the multimodal image color coded to the respective k-means cluster. Colors correspond to cluster identity 1-6. **f**, The stacked bar graph shows the simplex-normalized average pixel intensity’s channel contribution in each cluster. Color corresponds to discrete channel of the multimodal image stack. **g**, Regions of interest are selected along the cortex-medulla axis and a full SRS hyperspectral image sweep (512×512 pixels) of the C-H stretching region is obtained at the same time. The mean SRS spectra and 1-sigma error bands are plotted for each region. **h**, Phasor plots of all pixels in the ROIs separate spectra based on phase and amplitude. The plot is duplicated, and color coded according to region. Scale bar is 700 µm, corresponding to the large WSI in **a**. DM, Dichroic Mirror; EOM, Electro-Optic Modulator; OB, Objective Lens; CO, Oil Condenser; SF, Stokes Filter; PD, Photodiode; SRL, Stimulated Raman Loss; SRS, Stimulated Raman Scattering; TPF, Two Photon Fluorescence; SHG, Second Harmonic Generation

**Figure 2.**
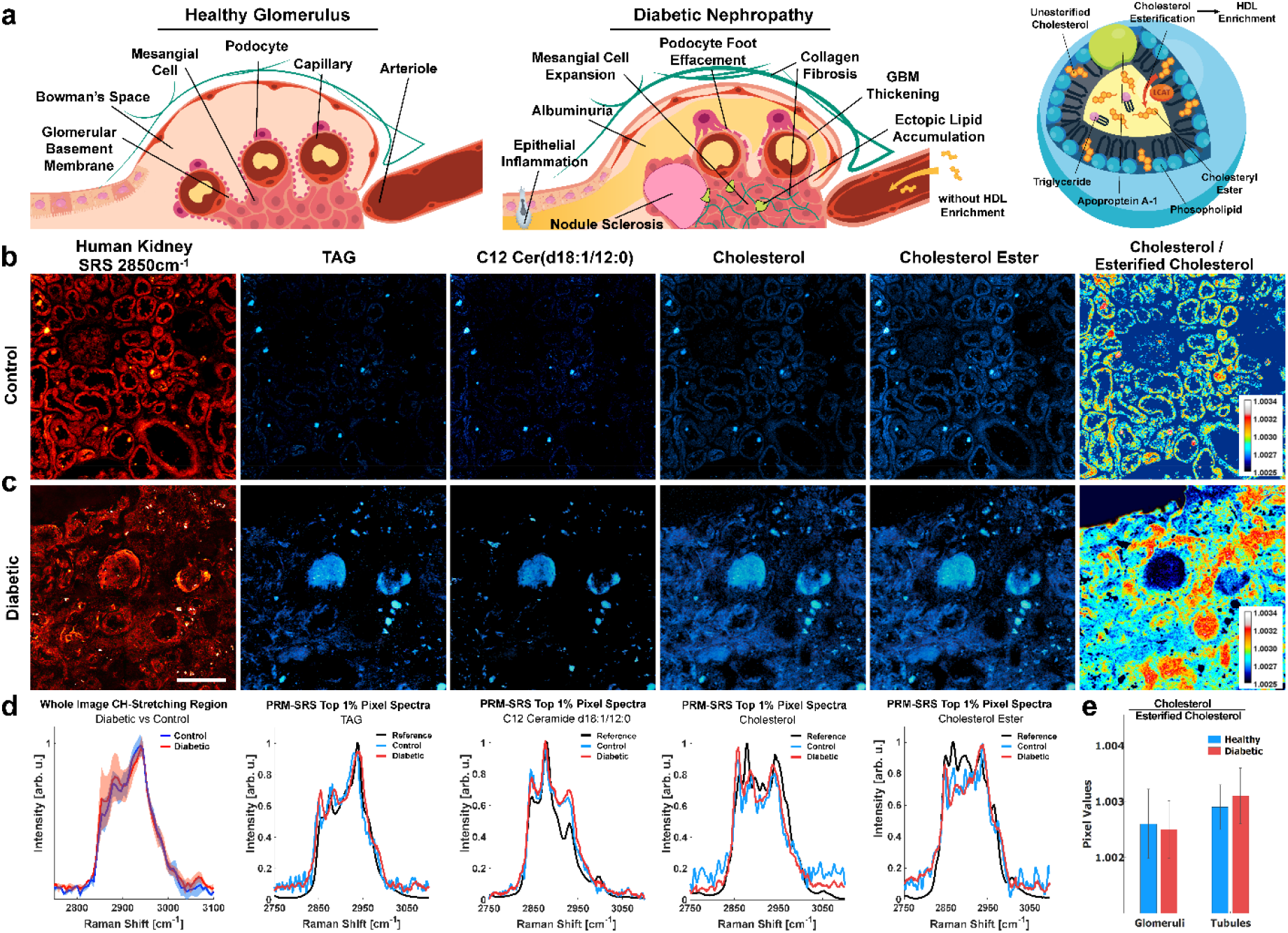
Label-free lipid subtype visualization in situ using PRM-SRS depicts cholesterol enrichment in diabetic kidney samples. **a**, Illustrated are some of the pathological features observed in the glomerulus in diabetic nephropathy compared to a healthy glomerulus and potential link with high free cholesterol and low serum HDL caused by lack of HDL enrichment. **b-c**, SRS images of human kidney at the C-H_2_ stretching peak (Left) and PRM-SRS images of reference-matched lipid subtypes of interest showing the spatial distribution of similarity scores at the same contrast level. **d**, HSI profile (Left) shows the Raman Spectra for all pixels in the HSI, with a red dashed line to indicate the 2850cm^−1^ image above. Spectra of the top 1% of similarity score pixels overlaid on the reference spectrum for each lipid subtype show confer PRM-SRS images. **e**, Ratiometric image of cholesterol and esterified cholesterol similarity scores show highest levels in eosinophilic bodies and within tubule epithelial cells, and not glomerular capillaries and podocytes. DKD tissues show higher relative free cholesterol in the tubulointerstital region, and lower relative free cholesterol in the glomeruli. This may be due to hyaline or protein in the glomerulus of DKD, leading to poorer reference matching of lipid subtypes in that region. Scale bar, 200 µm. Data presented are mean ± SD.

## Results and Discussion

### Unique Biomolecular Features of Diabetic Kidney Tissue

#### Alterations in Lipid Subtypes

Pathological variations in lipid subtypes, such as elevated cholesterol and an imbalance in quantities and qualities of sphingolipids, are often concomitant with other molecular and even morphological patterns in diabetic models^36–38^. Canonical biomarkers of diabetic nephropathy include mesangial expansion, nodular glomerulosclerosis, fibrotic extracellular matrix, ectopic lipid accumulation, and proteinuria (**Fig. 2a**). To examine changes of lipid subtypes within DKD tissue, we performed Penalized Reference Matching SRS (PRM-SRS) hyperspectral imaging (**Supplementary Fig. 3**). PRM is a fast and robust method for generating relative quantitation for a variety of user-defined standard components, such as lipid subtypes, by determining the angle or similarity score between two n-dimensional vectors^39^. These vectors can be discrete embeddings or continuous Raman spectra. In this case, the pixels in a hyperspectral image (HSI) captured by SRS each represent a chemical spectrum. The spectra were compared to reference spectra of lipid subtypes that are measured separately. Selected lipid subtypes, including TAG, C12 ceramide, cholesterol, and esterified cholesterol, in control and DKD kidney samples were compared. The total lipid image, ascribed to the C-H_2_ symmetric stretching Raman peak at 2850cm^−1^, is displayed alongside the similarity score images for these lipid subtypes in both control and diabetic tissues (**Fig. 2b-c)**. Pixel intensity corresponds to the similarity score, and diabetic core biopsies showed consistently higher similarity scores for each lipid subtype, indicating potentially higher relative concentrations of lipids. The spectra of pixels within the top 1% of similarity scores are plotted below the corresponding PRM-SRS images (**Fig. 2d**). While the normalized Raman spectra of control and diabetic tissues are very similar as a whole, the diabetic samples exhibit a larger standard deviation near the CH2 symmetric stretching region (2850cm-^1^). These findings suggest that diabetic samples may not only exhibit higher concentrations of select lipid subtypes, but that these discrepant lipidomic profiles may be spatially heterogeneous.

**Figure 3.**
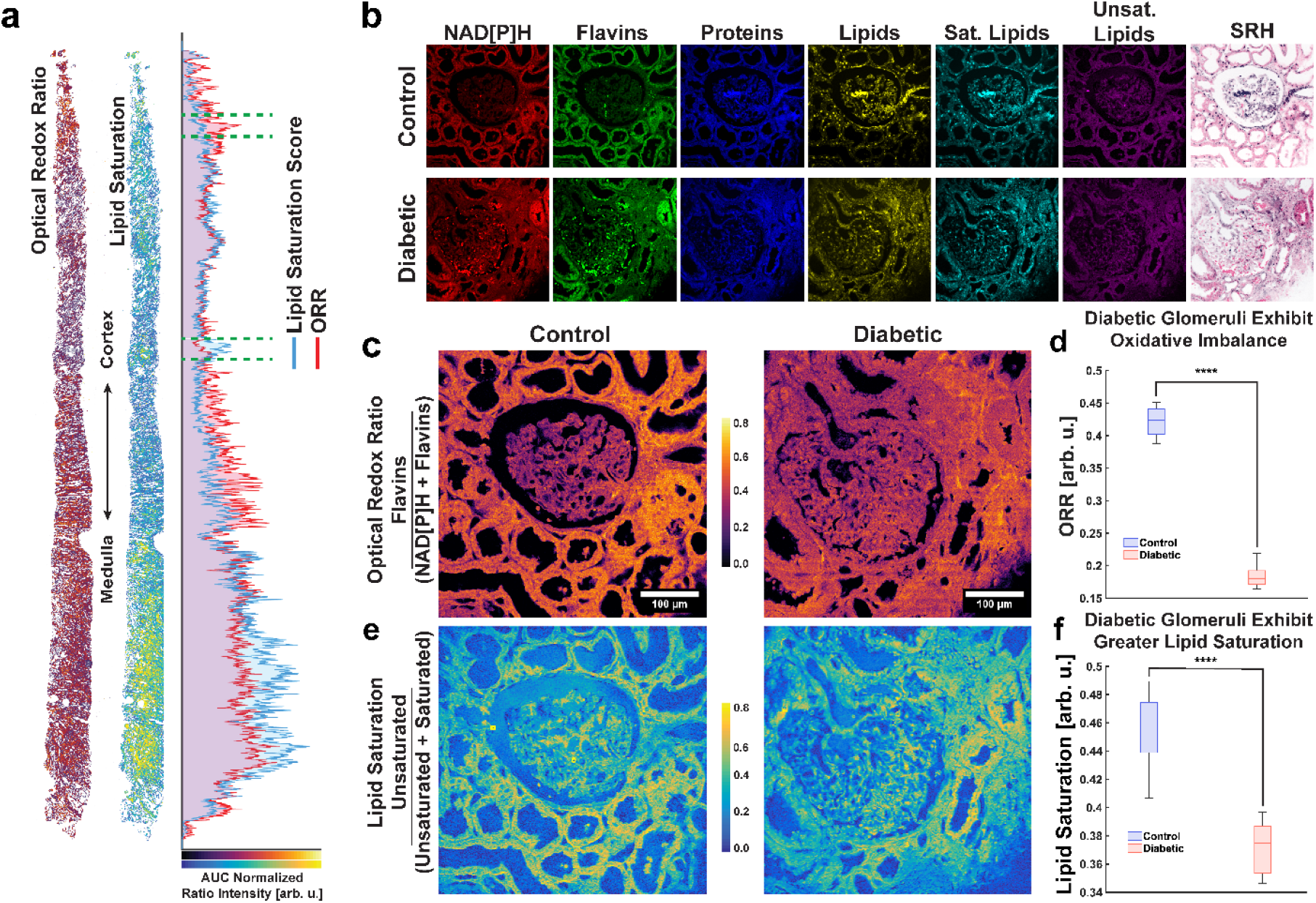
Diabetic glomeruli have an imbalance in redox ratio and lipid saturation. **a**, Core biopsies from diabetic patient along the cortical-medullary axis shows consistencies and differences between the optical redox ratio and degree of lipid saturation. Intensity profiles are plotted along this axis by ignoring background pixels and normalized by setting the areas under each curve equal to each other. **b**, Glomerular subregion images using SRS and TPF multimodal imaging, along with simple SRH processing. Note the mesangiolysis, marked by red blood cell stasis and regions of irregular matrix texture, in the diabetic glomerulus. **c**, Normalized optical redox ratio as a measure of oxidative stress and metabolism in healthy and diabetic glomeruli. **d**, Quantitative image analysis of ORR in glomeruli, n=9. **e**, Ratiometric images of lipid saturation in healthy and diabetic glomeruli. **f**, Quantitative image analysis of lipid saturation in glomeruli (n=9). Scale bar, 100 µm. Data presented are mean ± SD. \*\*\*\**P* < 0.0001.

Cholesteryl ester accumulation may be associated with cell proliferation and cancer aggressiveness^40^, but due to relatively low levels of fatty acid synthase proteins in human kidney tissues, the lipid profiles may be unique. We compared the ratio of cholesterol to its enriched esterified form by dividing the corresponding similarity score images and plotting the pixel values (**Fig. 2e**). These values may serve as an indicator for relative low-density lipoprotein (LDL) levels, and thus accumulation of free cholesterol^41^. Given reports of lipid droplet accumulation in diabetic kidney tissue^37,42,43^, it is reasonable to accept that slightly higher free cholesterol levels and lower enriched cholesteryl ester levels due to impaired cholesterol enrichment contribute to lipid accumulation in the kidney. Interestingly, we see that the glomerular and tubulointerstitial regions are inversely impacted in the diabetic sample with respect to the cholesterol esterification ratiometric result. Given the highly specialized roles of FTU’s such as the glomerulus, it is reasonable that these regions exhibit unique responses to dyslipidemia. Furthermore, glomeruli have been shown to participate in a complex network of activity within the kidney, and therefore underscore the necessity for spatial context^44^. These results warrant future studies to tie in functional discrepancies in diabetes, such as breakdown of filtration and reabsorption capacity, as well as the regional discrepancies in glomerular FTU’s.

#### Optical redox ratio and lipid saturation

Since differences in lipid subtype levels may also affect lipid metabolism within the tissue, we captured two metabolic indicators: the normalized optical redox ratio (ORR) and the lipid saturation^35,45^ The normalized optical redox ratio, defined herein as FAD/(NADH + FAD), can be obtained from TPF images of NADH and flavins such as riboflavins and FAD; and lipid saturation state, defined as unsaturated / (saturated + unsaturated), can be derived from SRS ratiometric images of saturated lipids at 2880 cm^−1^ and unsaturated lipid at 3011 cm^−1 46,47^. An example diabetic core biopsy showing both ratiometric image indicators is shown below, along with the average pixel intensity profile along its longitudinal axis (**Fig. 3a**), with glomerular regions of interest highlighted between the green dashed lines. Representative multimodal images are shown for both control and diabetic samples, along with stimulated Raman histology (SRH) images (**Fig. 3b**). The SRH images were generated by using two Raman wavenumbers at 2850 cm^−1^ (the CH_2_ stretch/lipid) and 2940 cm^−1^ (CH_3_ stretching/protein), respectively, according to previously published methods^48–50^. We observed both indicators are significantly lower in diabetic tissues (**Fig. 3c-f**). A lower normalized optical redox ratio may indicate less oxidation of fatty acids. A lower lipid saturation score suggests higher levels of saturated fats relatively to unsaturated fats, which can be typical of high fat diet and impaired HDL maturation. This is consistent with reports that increased ceramides, relatively stable fatty acids due to their higher degree of saturation, play a role in elevated reactive oxygen species (ROS) levels, and decreased desaturase (DEGS2)^37,51^.

In tissues such as the heart, liver, and kidneys, a lower redox ratio has been found to correlate with disease states such as diabetes. In diabetes, the reduced forms of pyridine nucleotides such as NADH and NADPH were elevated^52^. This appears counter-intuitive because a greater pool of NAD(P)H reductive potential suggests less oxidative stress, yet the opposite is observed. However, a shift toward utilizing anaerobic pathways to generate energy and thus away from oxidative phosphorylation could explain this paradox. Furthermore, It has been oberved that NADPH oxidase (NOX) activity is elevated in fibrosis showing a link between oxidative stress and fibrosis^53^. Likewise, a decrease in flavins such as riboflavins, flavin mononucleotide (FMN), flavin monooxygenase (FMO), and flavin adenine dinucleotide (FAD), could also account for a lower ORR. In diabetic retinopathy, for example, a flavin deficiency hampers the production of glutathione, a primary antioxidant defense^54^. These observations support our SRS methods are able to observe oxidative stress in diabetic glomeruli, as indicated by lower ORR and higher degree of lipid saturation.

### Morphological Features of DKD

#### 3D Imaging for Estimating Mesangial Fractional Volume in DKD

Metabolic alterations and the chemical microenvironment have a dynamic interplay, but prolonged dyslipidemia often results in irreversible structural damage^55,56^. One of the most widely used morphological analysis methods involves histopathological staining and inspection with light microscopy. Here we generated SRH images, which presents a more familiar semantic visualization of pathologies using multiple Raman features at once. Custom lookup tables (LUTs) were applied to these channels and blended in MATLAB to reveal subcellular morphologies in a remarkably similar manner to that seen in conventional H&E-stained images of tissue sections (**Fig. 4a-b**).

**Figure 4.**
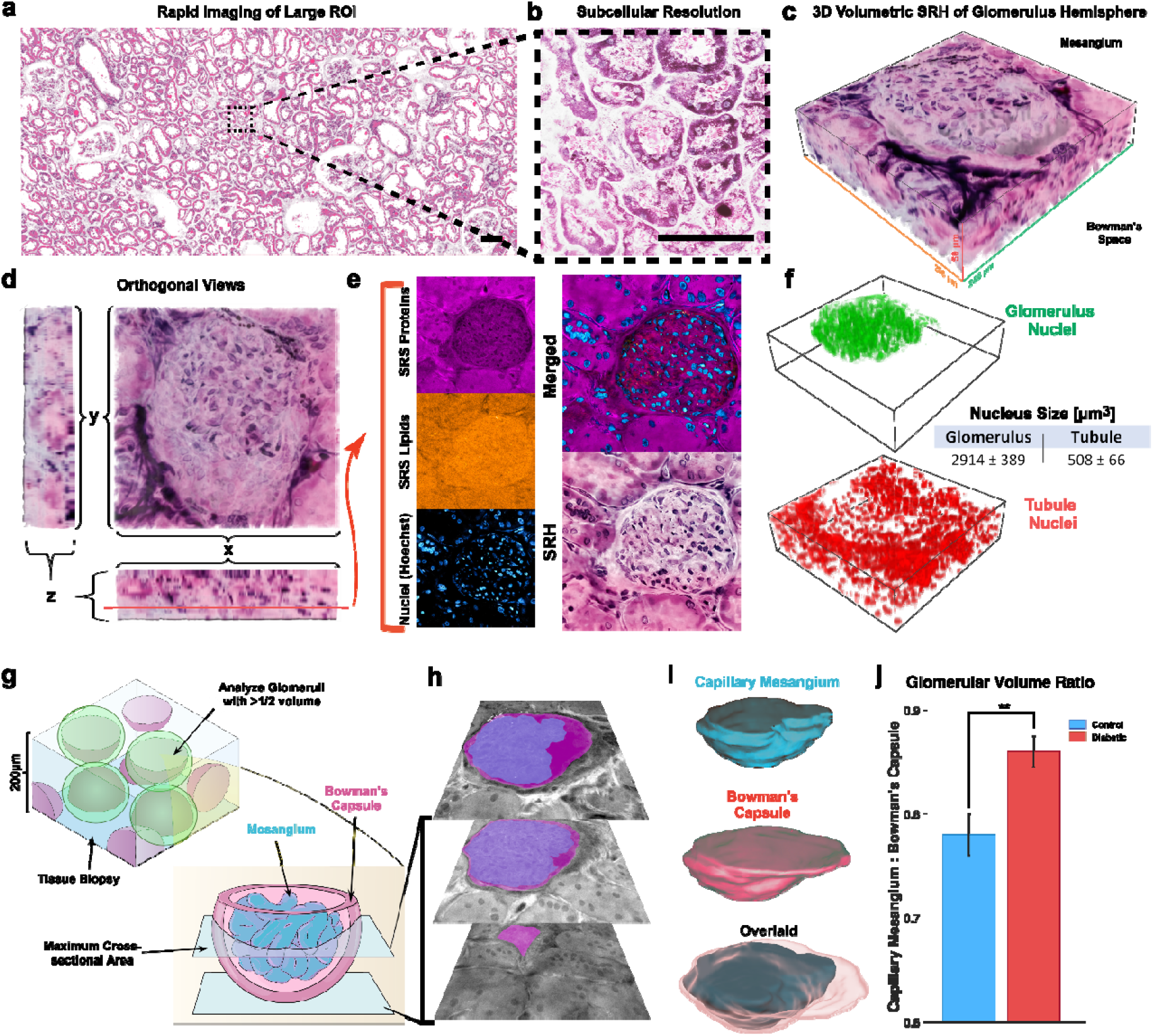
Label-free capabilities of SRH shows the rapid pseudo-coloring via linearly mixing protein and lipid SRS images of **(a)** large human kidney ROI and small subcellular resolution **(b)** in which nucleoli are visible. **(c)** Confocal scanning in the Z-direction can produce 3D pseudo-colored SRH images. **(d)** Orthogonal projections of (C). **(e)** SRS and TPF images of lipids, protein, and nuclei juxtaposed to the same image slice of the 3D stack shows that multiple visualization schema are possible. **(f)** Glomerular volumes can be segmented to inform the number of nuclei within the glomerulus and in the surrounding tubule cortex. Scale bar, 200 µm. **(g)** A schematic depicting the process of obtaining confocal image planes of glomeruli with at least a hemisphere within the imaging volume. Image planes between the maximum cross-sectional area and the vanishing point are retained. **(h)** Example of a few image planes of a glomerulus with the mesangium (cyan) and total glomerulus (magenta) overlaid on the CH_3_ stretching images. **(i)** The image planes are used to reconstruct volumes in ImageJ and MeshMixer. **(j)** Summary of glomerular factional volume [Vv(Mes/Glom)] between control samples and DKD samples shows that mean DKD glomeruli have significantly higher mesangial volume fractions compared to control specimens (n = 4 for each group). Data presented are mean ± SD. \*\**P* < 0.01.

This method was extended to 3D volumetric scanning of glomeruli to analyze the structure of these functional tissue units (**Fig. 4c-d**). While studies have delved into the capillary network of glomeruli using scanning electron microscopy (SEM)^57^, most reports examine these structures in 2D using light microscopy and PSI analysis. These methods underestimate mesangial volume by as much as 30% relative to 3D reconstruction methods^27^. Since SRS and TPF microscopies in our imaging platform are highly localized multiphoton processes, a 200 µm thick sample could acquire glomerular images at many focal planes by simply adjusting the height of the objective lens, as shown in **Fig. 4e**. Nuclei of glomeruli and tubules were segmented in 3D images and then counted (**Fig. 4f**). This enables quantitative assessment of cell number within subregions of FTUs such as the mesangium in disease states such as DKD.

Glomeruli are somewhat spherical tissue structures with a diameter of roughly 200 µm^58^. A 200 µm thick 3D image of a cortex sample might yield whole glomeruli, or more likely encompass several hemispheres or many medial cross sections as a 2D laminar slice with all glomeruli at the same focal plane. By optically sectioning 3D glomeruli, we capture more accurate morphological measurements and mitigate the impact of polar slices on statistical power (**Fig. 4g-h**). These slices were interpolated to a volume using ImageJ and MeshMixer (**Fig. 4i**). Only glomeruli with at least one hemisphere in the kidney biopsy volume, or with a distance of at least 100 µm between the largest and smallest cross-sectional area, were retained.

When estimating the glomerular tuft factional volume Vv(Mes/Glom), we are including the mesangium to include the mesangial cells and matrix, capillary endothelial cells, and podocytes, and consider the glomerulus to include the entirety of the renal corpuscle, including the Bowman’s space and the parietal epithelium. After identification and isolation of glomeruli, as described above, these glomerular and mesangial definitions allow for convenient morphological masking. Specifically, the entire glomerulus can be segmented using opening-closing structuring element operations, as well as size and shape filters described in the methods section below. The mesangium can be segmented by intersecting the glomerulus mask with the SRS image of protein to remove any bowman’s space background (**Supplementary Fig. S5**). The volume fraction is then a simple calculation of summing the number of pixels within each mask at each slice and dividing the result. We show that diabetic glomeruli have a significantly greater mesangial fractional volume than control, and support previous findings of underestimated volume fractions using 2D methods (**Fig. 4j**) ^22,23^.

The same samples were also analyzed by using a 2D single plane that mimics traditional methods, and by using each glomerulus’ maximal cross section within the volumes, respectively. Table 1 compares the mesangial fractional volume estimated using these two 2D methods and that using 3D hemisphere. 2D single plane measurements were calculated using the ratio of pixels in the mesangium to the entire glomerulus in a single plane view, like traditional methods. 2D max individual measurements were taken from a 3D image of each glomerulus, selecting for the plane with the largest cross-sectional area, and using this to calculate the ratio of pixels. 3D hemisphere measurements were taken from 3D images of glomeruli with either a defined polar and local maximum cross-section, or those with at least 100 µm contained within the 3D image volume.

**Table 1.** Comparison of Vv(Mes/Glom) measurement by different methods.

|  | 2D Single Plane |  | 2D Max Individual |  | 3D Hemispheres |  |
| --- | --- | --- | --- | --- | --- | --- |
|  | Control (n=20) | DKD (n=20) | Control (n=5) | DKD (n=5) | Control (n=5) | DKD (n=5) |
| **Mean Vv(Mes/Glom)** | **0.5482** | 0.5833 | **0.6344** | 0.7344 | **0.6167** | 0.6987 |
| **Standard Deviation** | **0.2138** | 0.2005 | **0.1783** | 0.0762 | **0.0450** | 0.0756 |
| **Standard Error** | **0.1073** | 0.0102 | **0.0844** | 0.0117 | **0.0016** | 0.0523 |

Interestingly, compared with control, the DKD samples exhibited a smaller variance between glomeruli in the 2D methods. This observation was in tandem with a lower incidence of polar cross sections showing a very small Vv(Mes/Glom). The lowest variance in measurements came from the 3D hemispherical method, which discarded several glomeruli based on the criteria previously described. Although volume measurements are more sensitive to small changes in radii than area measurements due to their cubed dimension, if only hemispheres are collected and analyzed, the variance within the population is mitigated, while the variance across populations, such as control and diabetic, can be comparatively more significant. However, greater precision may also come at the cost of longer imaging times even when restricted to glomerular FTU’s. Although 2D histology may be faster, they are susceptible to physical slicing deformations. More accurate glomerular measurements can be achieved using multiple sections, but this entails more sample preparation and co-registration^30^. For these reasons, optical sectioning via lightsheet fluorescence microscopy (LSFM) may be best suited for high throughput, contextual, glomerular measurements^44^.

#### Rapid Glomeruli Segmentation and Analysis using AI and SRH

Stimulated Raman Histology (SRH) has had a transformative impact on label-free imaging technology because it allows for tissue characterization using data from multiple Raman images in a familiar color scheme. To leverage histology’s seamless integration of spatial morphology and chemical specificity, we performed SRH on multiple control and diabetic patient samples and tested a popular convolutional neural network, called DenseNet, used in many artificial intelligence (AI) imaging applications. The goals were to determine whether label-free imaging affords the rapid identification of glomeruli, and whether glomeruli can be distinguished as being control or diabetic.

Using a fine-tuned SRH pipeline to generate realistic histological images for H&E and PAS (**Supplementary Fig. 6**), we tested DenseNet performance using the HALO AI platform (IndicaLabs). Both SRH and stained adjacent sections performed well in DenseNet segmentation of glomeruli (**Fig. 5a**), demonstrating that images collected with a label-free approach function just as well as ground-truth staining for this purpose. This is significant because glomeruli are major target FTU’s in the diagnosis of diabetic nephropathy, rapidly accomplishing this without the use of staining reagents can allow for multiplexed analysis on the same tissue slice.

**Figure 5.**
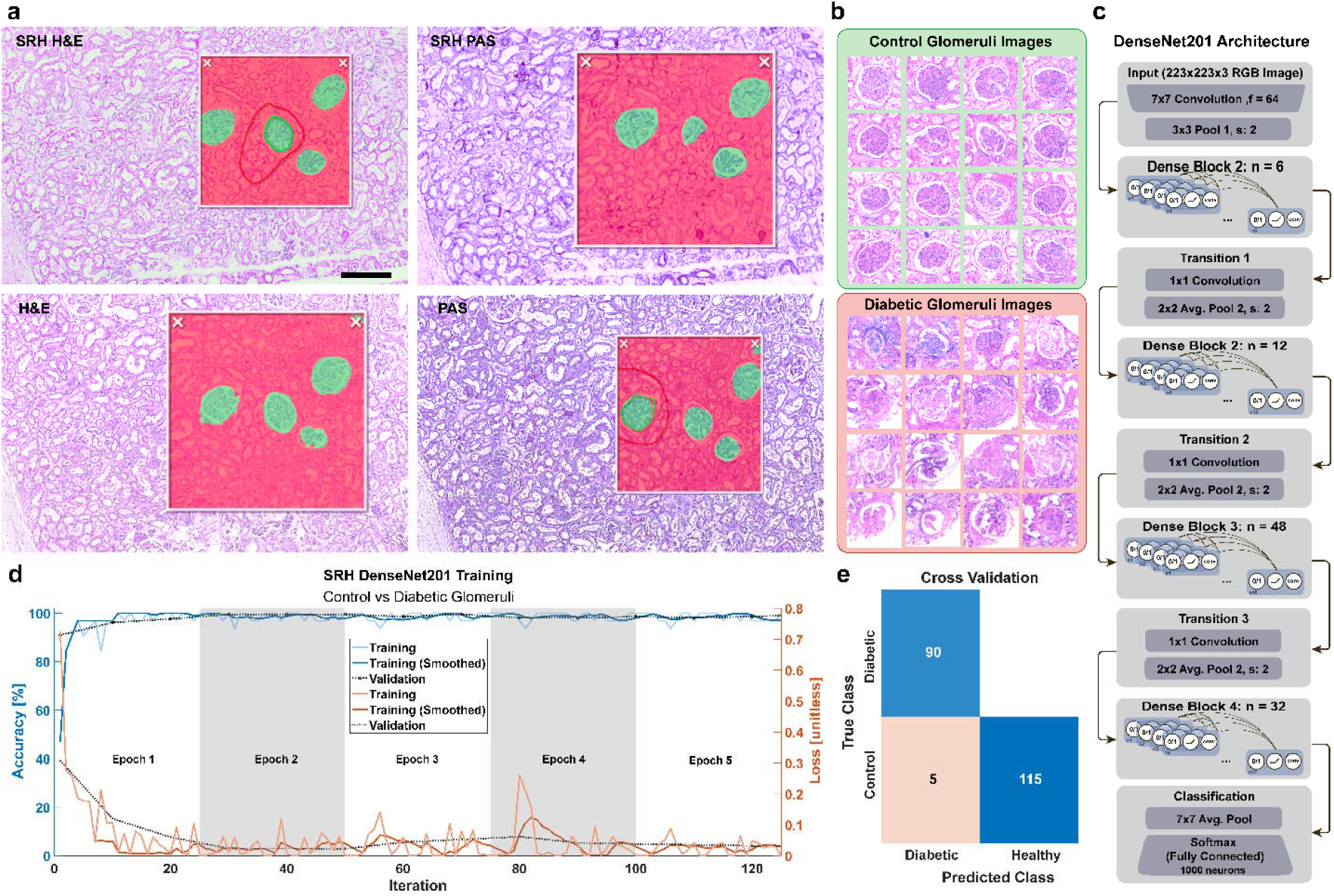
Neural Network detects and classifies glomeruli using only Stimulated Raman Histology. **a**, Adjacent kidney sections from the control group were imaged using label-free SRH and ground-truth histological staining for both H&E and PAS. Images were imported into the HALO AI platform by IndicaLabs and single glomeruli were manually annotated in each image. In all cases, SRH efficiently detected glomeruli just as well as histologically stained samples. **b**, 16 representative images of glomeruli were cropped from various patient samples in both control and diabetic groups. **c**, The network architecture of DenseNet201 is briefly explained using the MATLAB R2024a release implementation. **d**, The DenseNet201 network was trained using 5 Epochs, with 25 iterations per Epoch using a single Nvidia RTX 4080 and took less than 15 minutes. **e**, The cross validation matrix shows an accuracy of 0.9762, sensitivity of 1, specificity of 0.9583, and precision of 0.9474, resulting in an F1 score of 97.22%. Scale bar: 250 μm. H&E: Hematoxylin and Eosin, PAS: Periodic Acid Schiff, SRH: Stimulated Raman Histology,

**Figure 6.**
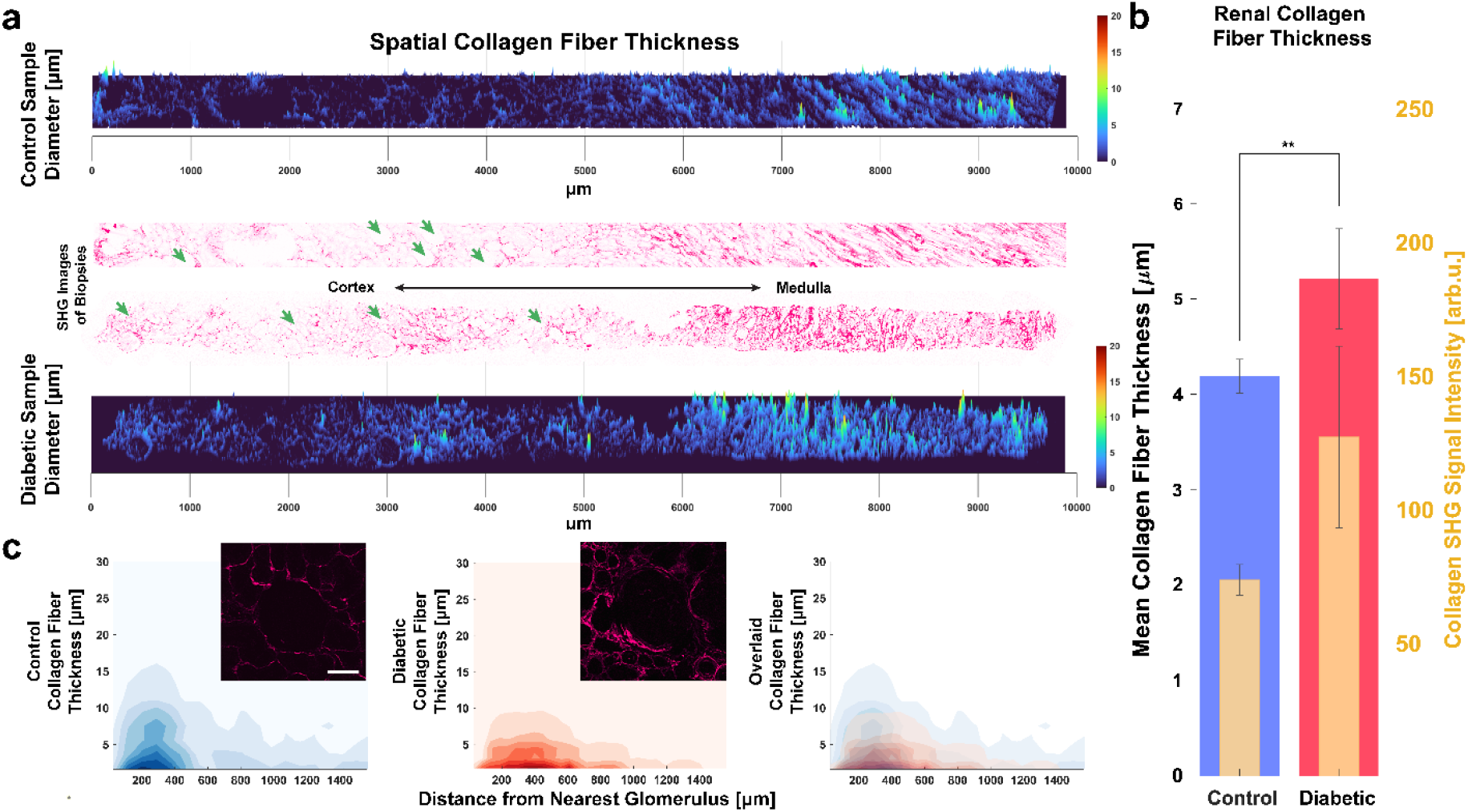
Spatial collagen analysis of human kidney biopsies using SHG imaging. **a**, Spatially-mapped fiber thickness of representative SHG images of control and diabetic biopsies along the cortical-medullary axis. Both samples have distinguishably thicker collagen fibers towards the medulla side, as well as peaks of fiber thicknesses in the cortex near glomeruli (marked with green arrows), vessels, and arteries. **b**, The mean fiber thickness of all fibers, as well as the mean SHG intensity for both control and diabetic samples show significantly higher levels in diabetic samples. n=4 tissue slides per group. **c**, 2D contour plots show the density of collagen fiber thickness measurements versus distance from the nearest glomerulus center in control and diabetic samples (n=20 glomeruli per group). Representative SHG images of glomeruli are shown to illustrate how fiber thickness tends to diminish further away from the glomerulus. Error bars represent one standard error of the mean. Data presented are mean ± SD. \*\**P* < 0.01.

After glomeruli were identified, they were automatically segmented (**Fig. 5b**) and fed into a training dataset for use in the DenseNet201 implementation in MATLAB (**Fig. 5c**). This network consists of Dense Blocks, comprised of composite functions that batch normalize, filter, and convolve features from the image, and Transition Blocks, comprised of a convolution and pooling layer to down-sample features for efficient computation. With only 5 Epochs of 25 iterations each, 80 diabetic glomeruli and 100 control glomeruli reached a classification accuracy of 98% (**Fig. 5d**). Using the 10% reserved set from each group, along with resampling of the original training dataset, glomeruli were distinguishable between control and diabetic origination with an accuracy of over 97% (**Fig. 5e**).

We employed DenseNet due to its common use, its underlying integration of residual network architecture robustness and the visual acuity of InceptionNet^59,60^. With off-the-shelf tools such as HALO AI and MATLAB, its implementation was straightforward. New tools are continually developed with the focus on meaningful histological feature extraction, such as HistoLens, which was demonstrated on renal histology images^61^. Furthermore, we felt that a neural network, which has an implicit goal of mimicking human interpretation, was the appropriate benchmarking tool for SRH, which has an implicit goal of recreating familiar color semantics.

#### Spatial Renal Fibrosis Analysis

Amount of renal fibrosis is a morphological indicator of chronic kidney disease and can be assessed using trichrome staining and fluorescent probes for both collagen and collagen binding protein^14^. Spatial analysis of collagen in-situ can be analyzed by area covered by collagen fluorescence and manually-measured fiber thicknesses using correlative SHG and TEM^62–65^. However other collagen characteristics that can provide insights into pathologic fibrosis include fiber orientation, strength, and relative density. Anisotropy analysis, also referred to as Histogram of Oriented Gradients (HOG) extraction, is one way of capturing these data^66,67^. We quantified type 1-3 collagen fiber angles from SHG images using the same Stokes beam (1031nm) from the SRS imaging. Without using any stains or other labels, we highlighted distinct regions of highly oriented fibers in the interstitium (**Supplementary Fig. 7a-b**). Here we binned the image into smaller areas to capture finer detail (**Supplementary Fig. 7c**), and thresholds for SHG intensity or anisotropy (**Supplementary Fig. 7d**). Anisotropy analysis separated kidney cortex from medulla and ensures that comparative regions are compared between control and diabetic samples.

To rapidly retrieve fiber thickness measurements throughout the sample, whole-slide SHG images were transformed into distance maps in which pixel intensity corresponds to distance to background (**Supplementary Fig. 8**). The regional maxima of this distance map correspond to the radii of collagen fibers. This measurement corroborates more manual metrics such as using the polar vector angles to orient perpendicular intensity profiles and capturing the full-width-half-maximum of the result. Plotting these measurements in a third dimension (also commonly known as a 2.5D image) shows that for both a diabetic and control tissue section along the cortical-medullary axis there are thicker collagen fiber diameters near the medulla side, as well as spikes in fiber thickness near glomeruli (highlighted by green arrows), vessels and arteries (**Fig 6a**). Utilizing this method to measure the whole tissue slide, we found that that the collagen fibers in the diabetic samples are nearly 25% thicker than the control samples, in addition to exhibiting higher SHG intensity (**Fig. 6b**). Interestingly, there was distinct local anisotropy near glomeruli especially in cases of glomerulosclerosis, vascular hyalinosis, and interstitial fibrosis. Separating the SHG images into glomerular subunits using Voronoi tessellations (**Supplementary Fig 5o**) allows us to plot the density of collagen fiber thickness measurements with respect to their distances from the center of the nearest glomerulus (**Fig. 6c**). Using anisotropy analysis, we automatically discount any medullary collagen that may be included near some glomeruli. Our results show that diabetic samples have thicker collagen fibers farther away from glomeruli.

This underscores the significance of spatial context in biology, as there have been recent strides in understanding the origin and progression of renal fibrosis^68–70^. Glomeruli and medullary thick ascending limbs are found throughout the tissue, and both are particularly susceptible to injury. Likewise, pericytes are found in both the cortex and medulla, and may be the most significant progenitors of collagen-producing myofibroblasts in the kidney^70^. Considering mesangial cells of glomeruli are known to be pericyte-like, it is prudent to know when and where these injuries take place, as well as finding dimensions that can discriminate diabetic pathologies.

## Conclusion

Our multimodal imaging and analyses methods enable visualizing morphological manifestations and responses to biomolecular changes at a subcellular level without destroying or wasting tissue. From a single microscopy setup, several analyses have been generated to illustrate both structural and biomolecular markers of control and diabetic kidney disease tissue. These include 3D quantification of glomerular mesangium, spatial collagen fiber thickness, optical redox ratio, and lipid saturation. Additionally, we leveraged a custom SRH pipeline and AI to automatically detect and classify glomeruli as either control or diabetic without using any staining reagents. We also demonstrated that this platform preserves the native orientation of the tissues by avoiding excessive physical slicing and allows for more representative volumetric data of the mesangial volume fraction.

This study is the first label-free multimodal imaging analysis of structural and molecular biomarkers in diabetic kidney disease, highlighting the potential of a single microscopy setup to dissect a multiplexed dataset while conserving tissue samples, and to inspire new research into the connection between dyslipidemia and kidney disease. Notably, we address preliminary results that indicate elevated lipid saturation and lower relative esterified cholesterol with respect to free cholesterol, as well as relatively higher NAD[P]H and lower flavin levels in diabetic samples. Potential future targets for analysis thus include mitochondrial dysfunction and transcriptional changes in co-factor enzyme production.

Future studies may iterate on these first demonstrations. For example, 3D digital histology may benefit from further segmentation of cell types, not just nuclei in certain regions. Additional stains can also improve diagnostic power and fully leverage the label-free hyperspectral imaging platform. These stains can be further improved using cycle generative adversarial networks (cycle GANs) to make these SRH images appear more standardized so that a database can be established^71^. It should be noted that SHG of collagen only visualizes type 1-3 collagen due to their non-centrosymmetric structure, and thickness measurements and SHG signal intensity depend on fiber orientation and position relative to the focal volume. Since collagen fibrosis is tightly regulated by lipid metabolism and oxidative stress, future studies are necessary to determine the extent to which the SHG intensity and collagen anisotropy may predict other nephropathies such as lipid subtype differences, normalized optical redox ratio, and mesangial fractional volume.

## Materials and Methods

### Kidney Biopsy Preparation

Human kidney samples were obtained from The Kidney Translational Research Center (KTRC) at the Washington University School of Medicine in St. Louis under an approved IRB protocol. Samples used for 2D imaging were fixed using 4% PFA or formalin and after washing were preserved as paraffin embeeded tissue blocks prior to sectioning at 7 microns. Samples used for 3D imaging were fresh frozen and stored at -80 degC in OCT until sliced with a sliding microtome (HM 450, Epredia) at 200 µm and cleared using 8M urea for 48 hours at room temperature. Samples were submerged in 1ug/mL Hoechst 33342 (Thermofisher) for 15 minutes to stain nuclei for co-localization verification. Samples were imaged between 1mm thick glass slides and number 1.5 thickness cover glass (Erie Scientific).

### 3D Microscopy

An upright laser-scanning microscope (DIY multiphoton, Olympus) with a 25x water objective (XLPLN, WMP2, 1.05 NA, Olympus) was applied for near-IR throughput. Synchronized pulsed pump beam (tunable 720–990 nm wavelength, 5–6 ps pulse width, and 80 MHz repetition rate) and Stokes (wavelength at 1032nm, 6 ps pulse width, and 80MHz repetition rate) were supplied by a picoEmerald system (Applied Physics & Electronics) and coupled into the microscope. The pump and Stokes beams were collected in transmission by a high NA oil condenser (1.4 NA). A high O.D. shortpass filter (950nm, Thorlabs) was used that would completely block the Stokes beam and transmit the pump beam only onto a Si photodiode for detecting the stimulated Raman loss signal. The output current from the photodiode was terminated, filtered, and demodulated in X with a zero phase shift by a lock-in amplifier (HF2LI, Zurich Instruments) at 20MHz. The demodulated signal was fed into the FV3000 software module FV-OSR (Olympus) to form the image during laser scanning. All SRS images were obtained with a pixel dwell time 40 µs and a time constant of 30 µs. Laser power incident on the sample is approximately 40mW.

Second Harmonic Generation (SHG) was used to capture type 1-3 collagen images. The 1031nm stokes laser described above, with 300mW and a dwell time of 10us per pixel, was used with 5-frame averaging. Backscattered SHG signals were filtered using 465nm filter.

NADH and Flavin autofluorescence images were captured using the 800nm pump beam with 350mW and a pixel dwell time of 10us/px with 5-frame averaging. Backscattered signals were filtered using a dual filter cube of 460nm and 515nm. All wide-view tile-stitching and confocal z-scanning was controlled by the Fluoview software (Olympus).

### Data Analysis

#### PRM-SRS Lipid Subtyping

PRM-SRS was conducted according to previously published methods^72^. Briefly, pure lipid subtypes such as those shown in supplementary figure S1 were spectroscopically analyzed using spontaneous Raman scattering. These spectra were used as reference spectra. SRS HSI had each pixel scored using cosine similarity in the CH stretching region at multiple offsets. Similarity scores were had a corresponding penalty that varied according to the offset to account for non-linear, lensing, or other non-standardized equipment effects. Resulting images contain pixels with simplex-normalized similarity scores, whose intensity corresponds to the relative similarity of that pixel’s spectrum to the reference spectrum. These relative intensities are interpreted as relative concentrations.

#### Stimulated Raman Histology

Stimulated Raman histology was performed following published protocol^73^ for linear RGB blending. Briefly, custom lookup tables were applied to the protein (2940cm^−1^) and lipid (2850cm^−1^) Raman images and blended in the RGB color space using either MATLAB or ImageJ. For clarity, RGB histogram specifications were applied to slightly adjust the final result^74^. For 3D representations, the opacity of white interstitium was achieved using a sigmoidal transfer function in the MATLAB VolumeViewer alphamap field. Fine-tuned SRH images were generated by analyzing ground-truth samples stained with reagents after hyperspectral SRS acquisition of the same samples. For H&E pseudocoloring, which is comprised of two main colors, a blue-purple nuclei stain (Hematoxylin), and a pink interstitial stain (Eosin). These colors can be easily segmented in the L*a*b* color space to generate ground truth masks. The protein (2940 cm^−1^) and lipid (2850 cm^−1^) SRS images from the hyperspectral stack are unmixed using previously published methods^75^. Of note, three Raman peaks of interest comprise the majority of the variance: 2850 cm^−1^, 2880 cm^−1^, and 2940 cm^−1^. Nuclei were segmented from the ratiometric result by thresholding pixels with the highest lipid to protein ratio. The nucleic mask is then de-speckled, closed, and hole-filled using standard ImageJ binary functions. Custom LUTs are applied to the unmixed protein and lipid images and blended together in the RGB space in a similar fashion to other studies ^73,76^. To ensure the nuclei and background have enough contrast, the pixels within these masks have their L* decreased and increased, respectively, and their b* chromacity slightly decreased. Finally, RGB histogram normalization may be applied to ensure clarity and consistency ^74^. For the PAS SRH implementation, the stained ground truth images were preprocessed by enhancing contrast (raising the lower threshold by 10% and decreasing the upper threshold by 10%), and increasing the blue channel’s chroma and saturation by 10% each. This is done to provide a greater separation between the nuclei counterstain and the PAS aldehyde chroma. Once the image is converted to the L*a*b* color space, the PAS can be readily analyzed. Unlike the Eosin, which stains almost everything the same color, and can then be scaled in brightness proportionally with the Raman intensity, the hue and chroma of the PAS stain is more of a gradient. Period acid first oxidizes glycol groups found on saccharides in glycoproteins and mucin such as in basement membranes. The aldehyde product can then bind with Schiff’s reagent, and upon the release of a sulfonic group in washing steps, a reddish hue develops. The chroma of this hue is redder where the concentration of Schiff’s reagent is higher, however it unfortunately doesn’t scale linearly with Raman intensity. In the L*a*b* space, the areas with higher PAS staining correlate logarithmically with the difference between a* and b* chroma. It should be noted that other color spaces such as Hue Saturation Value (HSV) may also be used, however the L*a*b* space was implemented because Cartesian coordinates are easily interpretable with Raman intensity, and the histological stains employed are well suited to the a* and b* chromaticity representation. These axes measure green-magenta, and blue-yellow opponent colors, respectively. The chroma becomes redder with positive increasing a*, and bluer with negative decreasing b*. For images with another color gamut, a different segmentation method may be necessary. Unlike the more common RGB and CMYK color-spaces, the L*a*b* color-space is device independent. This offers advantages in color normalization methods to standardize datasets and establish uniform stain-augmentation in the future – a critical consideration for co-registration of spatial molecular imaging modalities.

#### Morphological Segmentation of Glomeruli from SHG

SHG images were analyzed using MATLAB. The collagen SHG signal is first morphologically filtered using a top-hat background subtraction with a disk structuring element of radius at least twice the largest visible fiber diameter. The result is median filtered to smooth small gaps, and then both dilated (to serve as a gradient mask) and inverted. The inverted image is eroded and reconstructed and is then used to find objects using regional maxima, corresponding to glomeruli and tubules. The resulting image undergoes a series of opening and closing to solidify the objects. An optional iteration can be applied to detect any missing objects by merging the result with the gradient image and repeating the last step. Finally, the object masks are dilated until they reach the gradient wireframe or intersect with another object mask. Code demonstrating this process is available on the Shi Lab Github account.

#### 3D Glomerular Tuft Volume Fraction

The CH_3_ Raman shift (2940cm^−1^) was used for the 3D image analysis of glomeruli because of its high intensity and optimal contrast. Glomeruli were manually identified and the planes between their maximal cross-sectional area and vanishing point were analyzed. The thickness of the glomerulus is the distance between those focus planes. The radius of the glomerulus is the average distance from the centroid of the glomerulus to the edge of the outer layer of the Bowman’s capsule at the plane with the maximum cross-sectional area. If no maximum cross-sectional area could be defined (i.e. the plane with the largest area occurs at the first or last plane of the 3D image) then the radius is assumed to be 100 microns. Those glomeruli that showed thicknesses of less than 90 percent of their respective radii were discarded. These may have been due to the region being a polar slice (less than a hemisphere), or the glomerulus not fully contained within the imaging volume, respectively.

Auxiliary analysis using the maximum cross-sectional area plane for the volume fraction estimation was also performed. For these measurements, only glomeruli clipped at the edges of the image were discarded, just as with the traditional 2D method.

#### Glomeruli Segmentation

Single plane hyperspectral SRS images of human kidney were acquired along with the collagen SHG images. The SRS hyperspectral image was projected to 2D using maximum intensity projection (MIP) for ease of viewing. The collagen SHG image was transformed into a mask by maximizing the contrast and closing gaps manually using the pen tool in ImageJ. The muscle MorphoLibJ plugin was used to generate a segmentation of the collagen mask, which was then intersected with the MIP image and thresholded to remove the empty spaces such as those between tubules. Pixels within the segmented regions correspond to a datum, which were then k-means clustered initialized with 3 clusters. Segments were pseudo-colored by cluster identity using lookup tables for visualization. Glomeruli were also segmented using HALO AI (IndicaLabs) using a DenseNet 2.0 model. After importing the stained and SRH images, manual segmentation of example glomeruli and tubulointerstitium are drawn. Then, a bounding box is drawn to indicate the analytical region containing both glomeruli and cortex tubulointerstitium to be analyzed. Green and red masks appear within this bounding box to indicate structures of interest.

#### Collagen Anisotropy

Collagen fiber direction and anisotropy, along with total collagen fiber signal, was extracted into 80um x 80um regional bins using a bespoke MATLAB code. Details of the methods are published elsewhere ^77^. In brief, the method applies a 2D discrete Fourier transform (DFT)– based algorithm to the SHG images and features a novel integrated periodicity plus smooth image decomposition to correct DFT edge discontinuity artefacts, minimizing the loss of peripheral image information that limits more commonly used DFT methods. Collagen fiber thickness measurements were achieved by transforming whole-slide SHG images into distance maps in which pixel intensity values correspond to its distance to the nearest background pixel using the bwdist function in MATLAB. Local Maxima were recorded and multiplied by 2, since the pixel distance is half the thickness. Background subtraction was done using a flatfield correction and top-hat filter in MATLAB with a disk morphological structuring element.

#### Statistical Analysis

A total of 2 DKD samples, and 3 control samples were used in this study. For lipid subtype, collagen anisotropy, and optical redox ratio calculations, 3 regions of interest were obtained from each sample. For glomerular volume fraction calculations, 4 representative hemispherical glomeruli were used in each group. A simple student’s t-test was used for all measures of significance.

## Funding and Acknowledgements

We are grateful for the support of the Washington University Kidney Translational Research Center (KTRC) for kidney samples and the HuBMAP grant U54HL145608. We also thank Amanda Knoten and Kristy Conlon for patient enrollment and sample preparation. We also thank the support of the NIH R21NS125395 and Sloan research fellow award.

## Notes

### Competing Interest Statement

The authors have declared no competing interest.

