## Supplemental File for "Label-Free Optical Biopsy Reveals Biomolecular and Morphological Features of Diabetic Kidney Tissue in 2D and 3D"

**Supplementary Information**

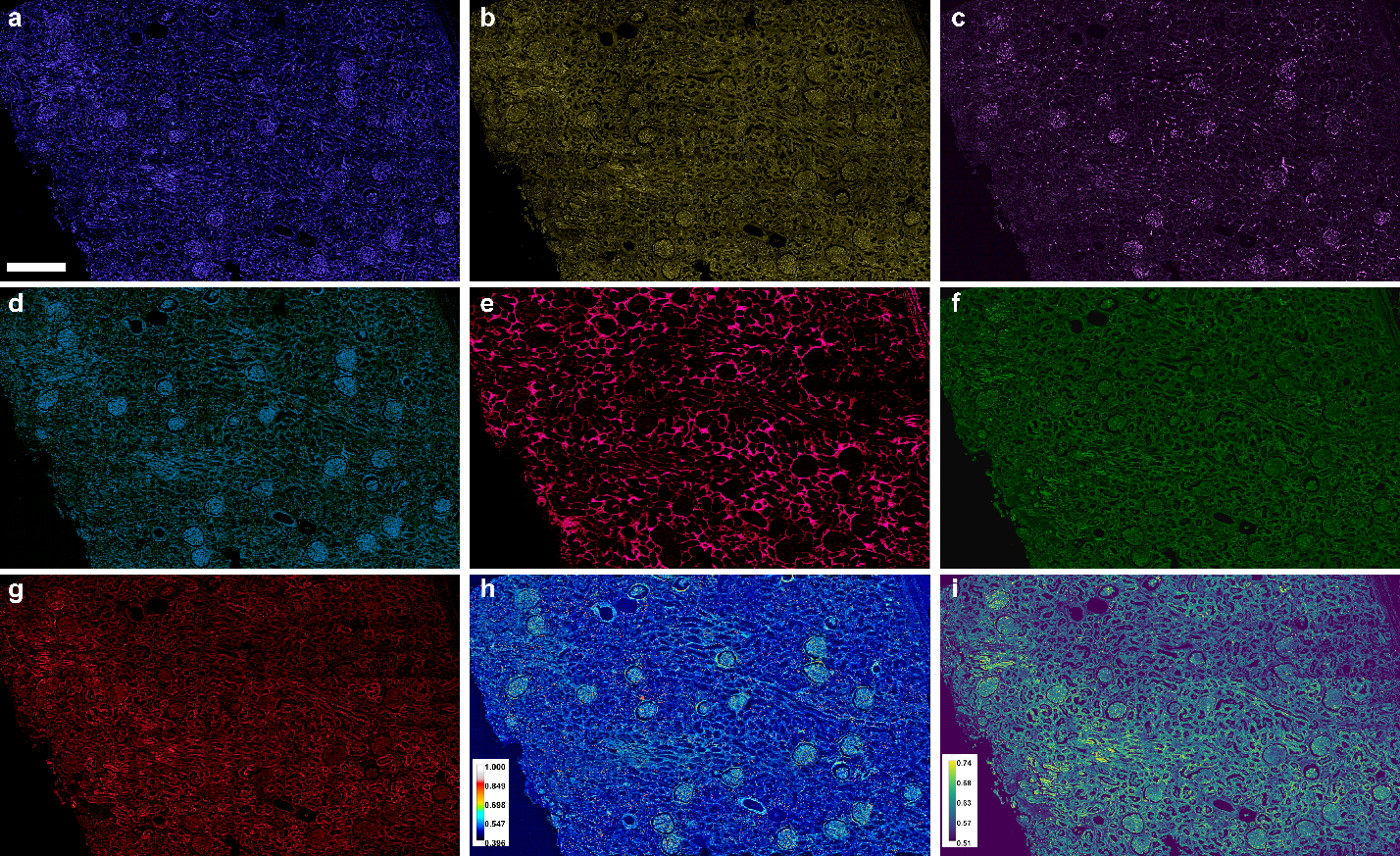

**Supplementary Fig. 1: Example multimodal images of control human kidney tissue. a,** SRS image of human kidney necropsy captured at the 2940 cm^-1^ Raman peak ascribed to protein. **b,** SRS image of the 2850 cm^-1^ Raman peak ascribed to lipids. **c,** SRS image of the 2880 cm^-1^ Raman peak ascribed to saturated lipids. **d,** SRS image of the 3010 cm^-1^ Raman peak ascribed to unsaturated lipids. **e,** SHG image captured at 1031nm/515nm ascribed to fibrillar collagen types 1-3. **f,** TPF image of flavins captured at 860nm/525nnm. **g,** TPF image of NAD[P]H captured at 780nm/460nm. **h,** Ratiometric image of lipid saturation calculated by the normalized ratio of d and c. **i,** Normalized optical redox ratio image calculated by the normalized ratio of f and g. Scale bar, 500 µm.

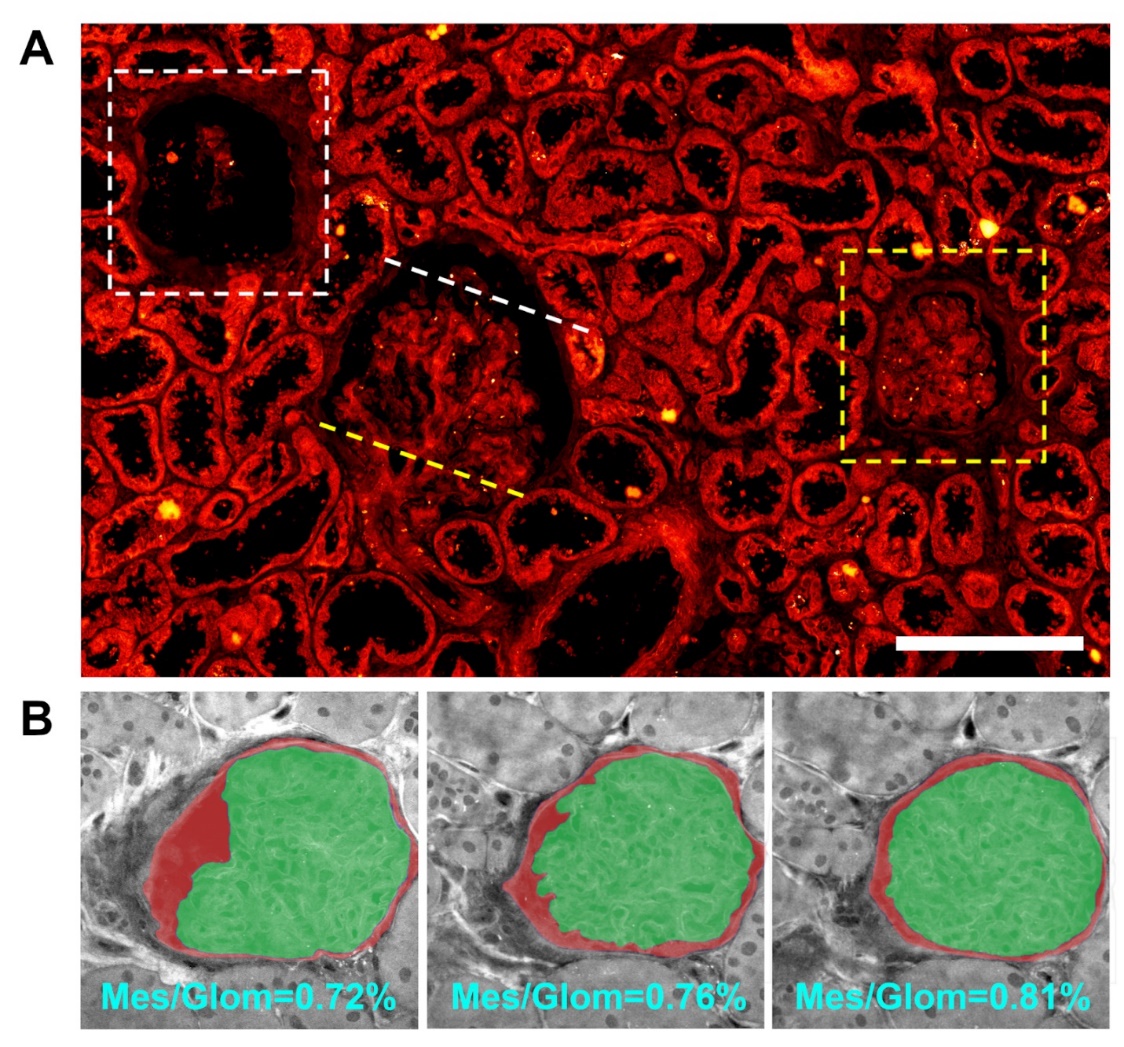

**Supplementary Fig. 2:** **2D human kidney biopsy shows dramatic differences in glomerular profiles**. **(A)** Glomeruli in various focus planes. White dashed box shows a glomerulus that was perhaps sliced distal to the arteriole, indicated by the white dashed line on the glomeruli sliced near the center. Yellow dashed box shows glomerulus that was perhaps sliced proximally to the arterioles, indicated by the yellow dashed line. **(B)** Focal planes of a single glomerulus only 15 µm apart from each other show significant differences in the area fraction of the mesangium (highlighted in green) in the glomerulus (highlighted in red). Scale bar, 200 µm.

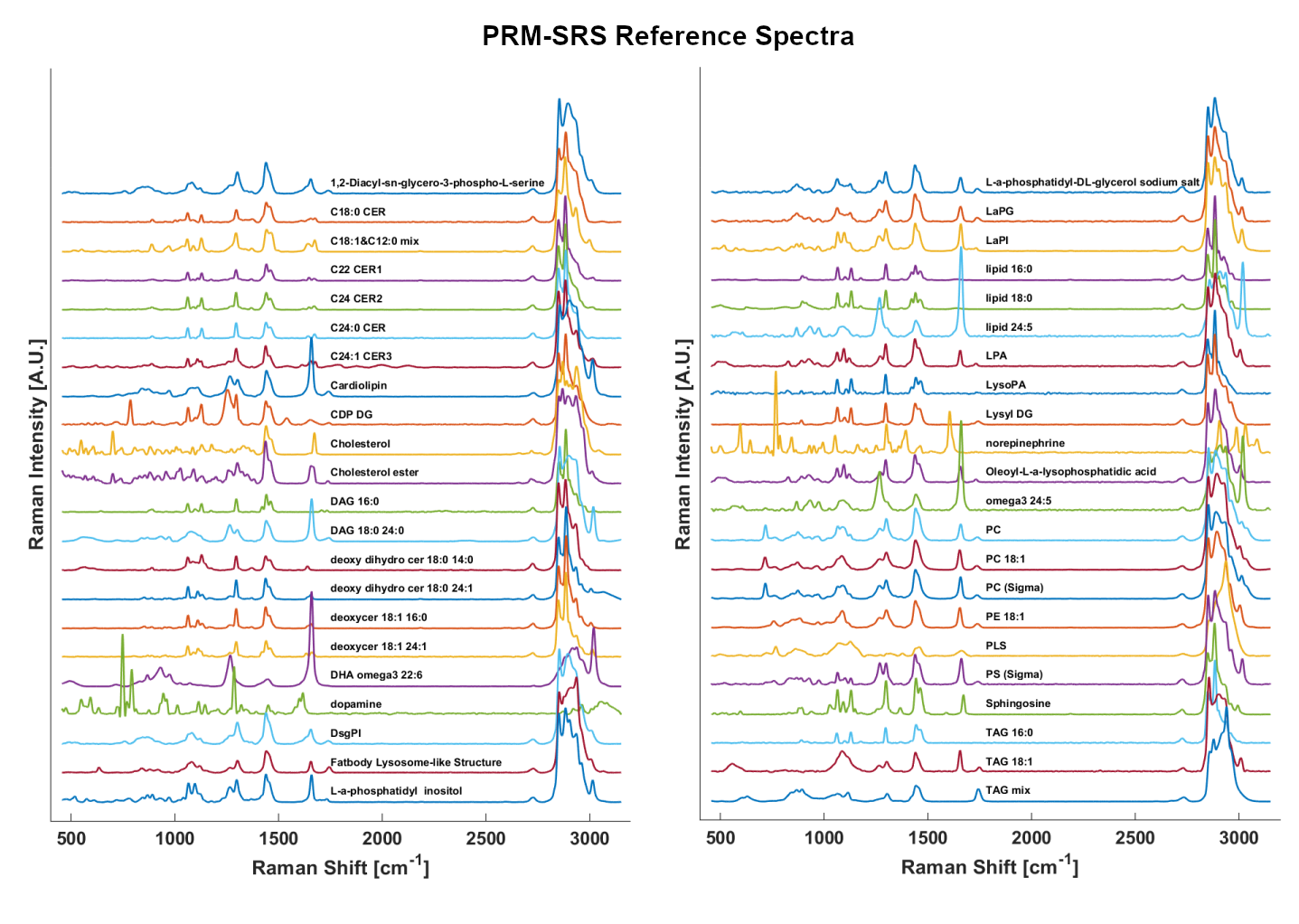

**Supplementary Fig. 3:** Spontaneous Raman spectra of lipid subtype standards used in PRM-SRS.

Each colored area in **Fig. 4c** is a datum that can be classified using k-means initialized with 3 clusters. These clusters were segmented and overlaid, along with the collagen, in **Fig. 4d**. The spectral profiles of glomeruli and the rest of the tubulointerstitium (**Fig. 4e-f**) show that for both control and diabetic samples, there is generally a higher protein to lipid ratio in both control and diabetic glomeruli. This reiterates that little sample preparation and signal processing is required for such a hyperspectral image. A large DKD kidney sample that was processed in this manner (**Fig. 4g**) illustrates some broad morphological differences such as a denser tubulointerstital collagen network (green), tubules (red) with less space in their centers, and glomeruli (cyan) with a higher mesangial area fraction.

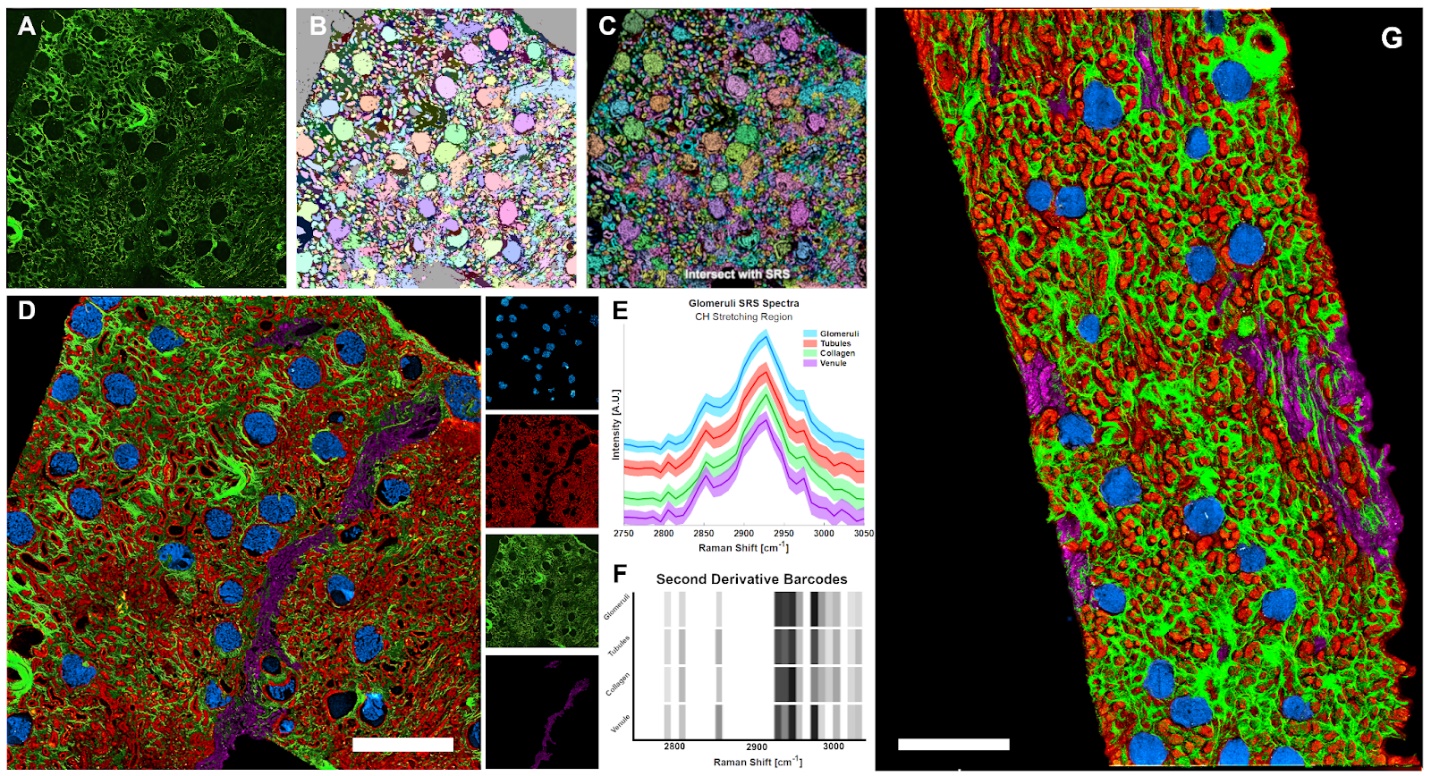

**Supplementary Fig. 4:** SRS hyperspectral imaging clusters morphologies **(A)** collagen SHG, used as a segmentation mask, to separate glomeruli/tubules with a watershed segmentation shown in **(B)**. **(C)** The resulting mask in (B) is intersected with the maximum intensity projection of the SRS HSI to remove background pixels and is then loaded into MATLAB. **(D)** Overlaid and separated pseudo-colored morphologies base on K-means clustering initialized with 4 clusters, with the pixels covered by collagen SHG (A) included as well. **(E)** A plot of the mean and standard deviation of the pixel spectra from each labeled cluster. **(F)** A second derivative barcode improves the visualization of the wavenumbers of interest in (E). **(G)** Diabetic Kidney sample analyzed with the same method reveals a much denser collagen intertubular network and inflamed tubule epithelial cells. Scale bar, 600 µm.

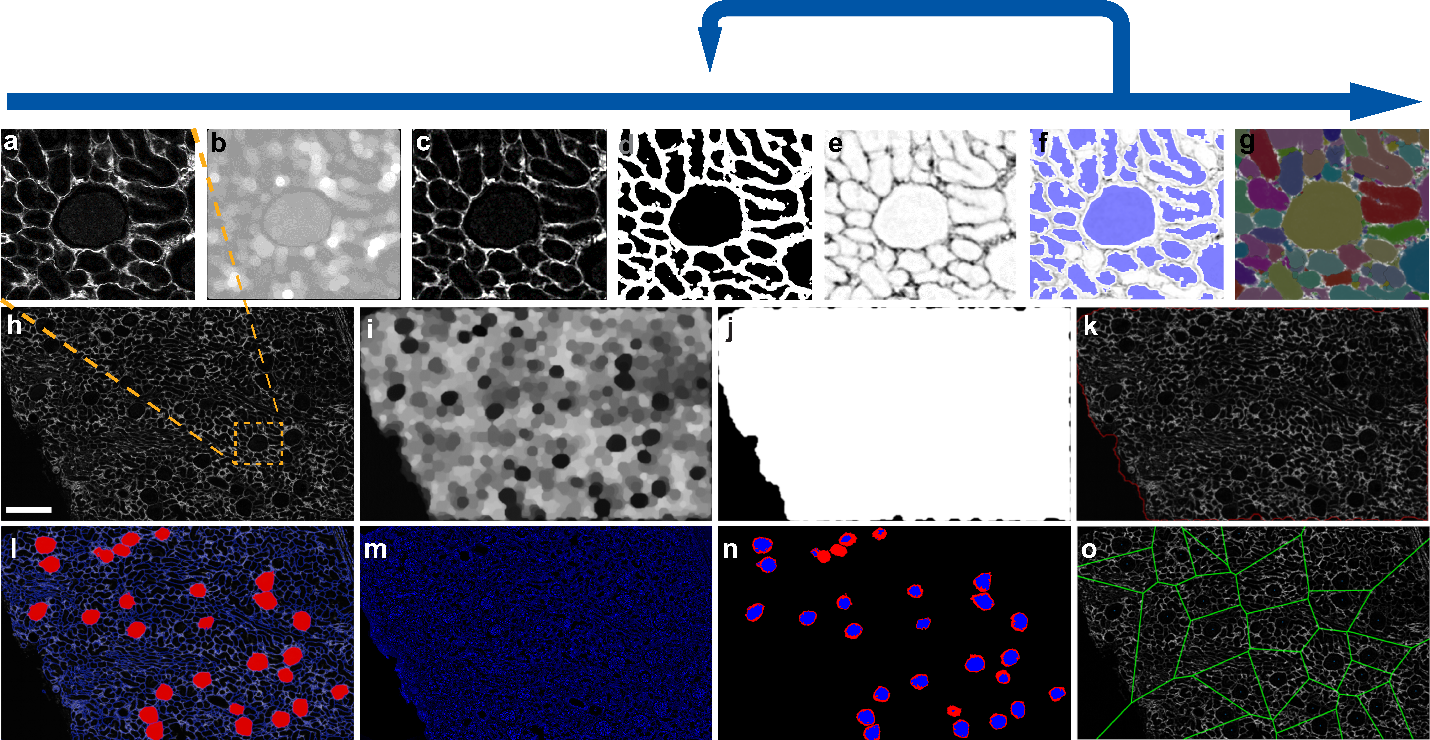

**Supplementary Fig. 5:** Targeted analysis of glomerular subunits using morphological filtering. **a,** Glomerular ROI enlarged from a large SHG image as an example. **b,** Top-hat background to be subtracted from the original SHG image. **c,** The result of an unsharp mask applied to the result of image a – image b **d,** The result of image c has small gaps bridged and dilated to generated a gradient magnitude segmentation function.  **e,** Image c is inverted to initialize the structuring element opening step. The regions of interest start at the peaks of this inverted image and erode the image to where the gradient intersects the fiber boundaries. **f,** Closing by reconstruction flattens maxima inside the regions of interest and highlighted in blue using the imregionalmax() and labeloverlay() functions in MATLAB **g,** Gaps or missing regions can be improved by iterating through steps d and f after intersecting the result from image f and image d. Finally labeled regions of interest are dilated until their borders intersect. **h,** A cortical region of a human kidney necropsy imaged using SHG microscopy. **i,** Following the opening and closing by construction steps above, a textured filter is generated. **j,** The textured image can be binarized and have its holes filled to create a mask of the tissue from the background or even the cortex from the medulla. **k,** The original SHG image of the kidney cortex is outlined in red following the mask generated from image j. **l,** Image i can also be used isolate glomeruli using morphological filters such as size (greater than 30,000 square microns) and circularity (greater than 0.8). **m,** A co-registered SRS image of the protein peak (2940cm^-1^) **n,** Glomeruli relative to the Bowman’s space is analyzed by intersecting the protein image m and the glomerular subunits from image l and binarizing the remaining protein signal and filling holes. **o,** Voronoi tessellations are generated by finding the centroids of the segmented glomeruli for further spatial analysis. Scale bar = 500µm.

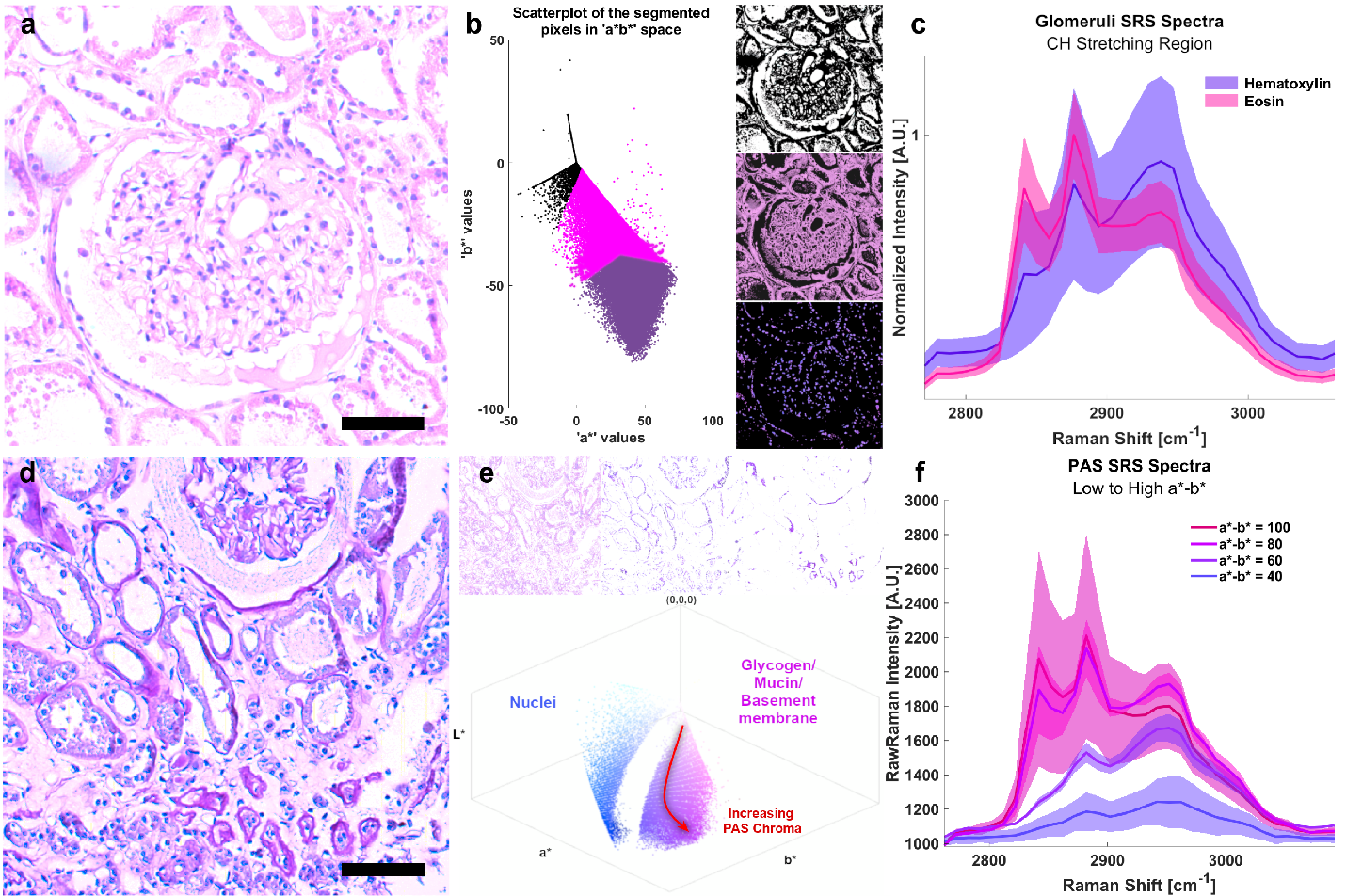

**Supplementary Fig. 6:** Stimulated Raman Histology (SRH) platform shows **(A)** H&E stained ground truth image from light microscopy post-SRS hyperspectral imaging. (B) The ground truth image is transformed into the L*a*b* color space and segmented to isolate background, eosin, and hematoxylin, and used as masks to capture the Raman spectra of those pixels shown in **(C)** with 1-sigma error bands**.** The nuclei stained by hematoxylin have a distinctly higher protein to lipid peak ratio so spectra were displayed after min-max normalization. (D) PAS stained ground truth image from light microscopy post-SRS hyperspectral imaging. (E) Ground truth image is preprocessed by enhancing contrast and increasing chroma of the blue colors to provide greater separation between the nucleic counterstain and the progressive PAS aldehyde chroma. The image was then transformed to the L*a*b* color space. The mucin stain becomes a gradient, with logarithmically increasing mucin chroma along the a*-b* axis. Snapshots of the L*a*b* thresholded image along the red arrow are shown above. **(F)** Several groups of pixels with indicated a*-b* quantities are plotted with 1-sigma error bands.

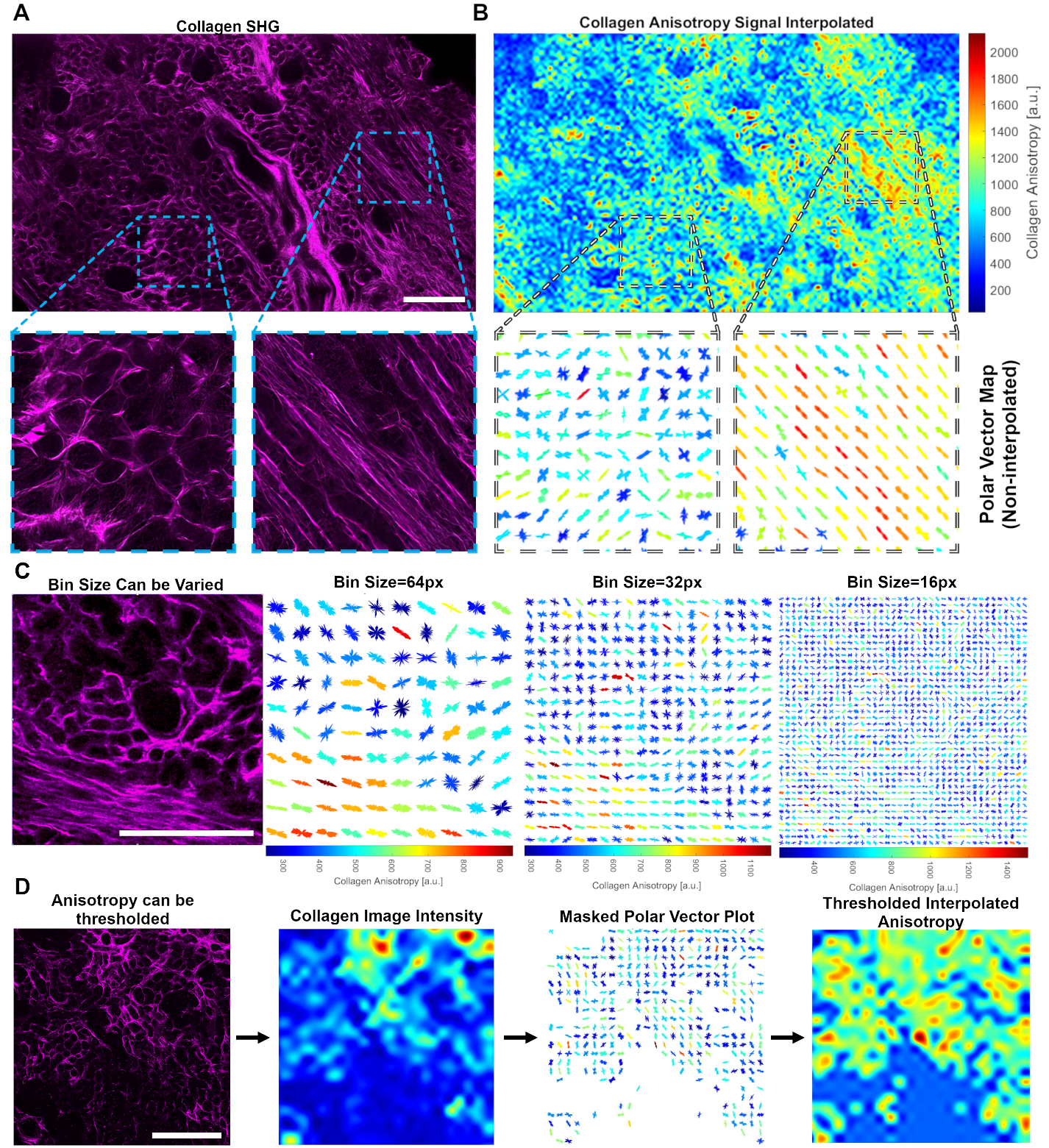

**Supplementary Fig. 7:** Fourier-based analysis of renal SHG images **A)** SHG image of collagen in a 50 micron-thick human kidney sample using a 1031nm stokes laser. **B)** Collagen fiber orientation represented as polar vector plots (plot separation: 80 µm x 80 µm). Plot shapes reveal preferential directions of fibers, whereas colors reflect anisotropy scaling (warmer colors = higher anisotropy) **C)** Bin sizes can be varied to capture polar vector representations of different portions of tissue. **D)** Anisotropy can be thresholded using either the collagen signal intensity (depicted) or polar vector length to eliminate unwanted signal. **E)** Collagen fiber thickness can be measured from an SHG image (i) as a whole by selecting the local maxima of a distance map (ii-iii). DN samples had a mean collagen fiber thickness 24.4% greater than the control samples (iv). Scale bar, 500 µm.

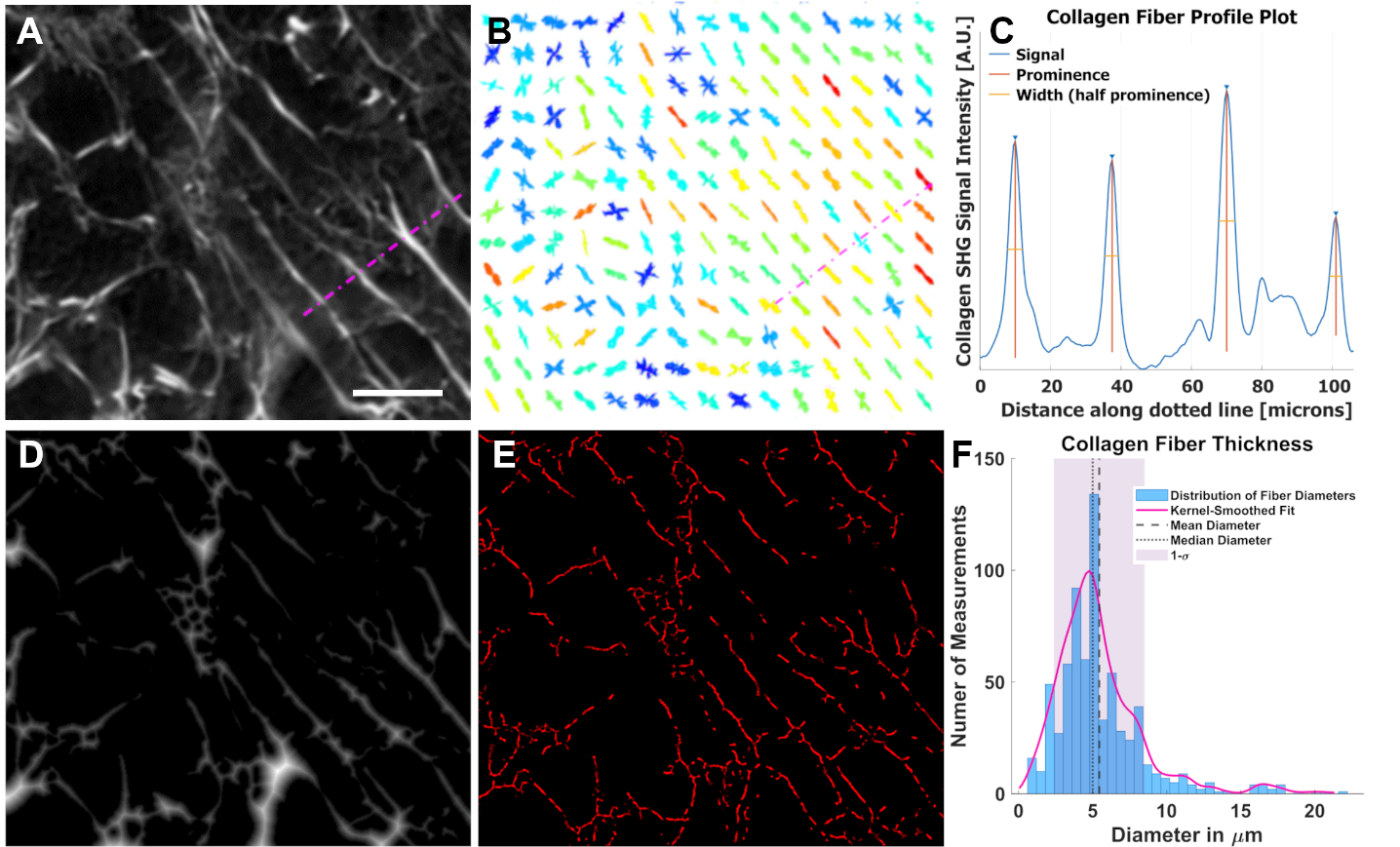

**Supplementary Fig. 8:** Collagen fiber thickness measurement strategy validation. **(A)** SHG image of human kidney captured at 1031nm/515nm shown in Figure 6E. **(B)** Anisotropy plot showing the polar vector angles of the same region using 16px bin sizes. A dotted line is drawn perpendicularly to the average vector angle that it crosses in both A and B. **(C)** The intensity profile is plotted along the dotted line drawn in A and B, with widths at half prominence around 4.5 microns. **(D)** The distance map converted image of A, also shown in Figure 6E, undergoes regional maxima filtering shown in **(E)** to select pixels corresponding to collagen fiber radii. **(F)** the pixel values of D are multiplied by 2 and plotted in a histogram. The average diameter is 5.4 microns.
